# Study on Changes in Tourism Land and Influence Factors in Mountain Areas: A Case Study of Luanchuan County, China

**DOI:** 10.1101/2021.03.03.433724

**Authors:** Yanna Xie, Qingxiao Zhu

## Abstract

The rapid development of the tourism industry in mountain areas has resulted in intense changes in land use structure and exerts a strong influence on terrestrial ecosystems. This paper takes Luanchuan County (typical mountainous terrain in western Henan Province, China) as an example and employs land use data from different times and spaces and the binary logistic method to study tourism land spatial variation and influence factors in mountain areas. The research shows that: (1) spatial variation in land use in scenic spots reveals a chain reaction of land use type transformation caused by construction land expansion, a key driving force of spatial variation in land use and land use type transformation, and (2) the changes in tourism land use types result from human utilization and development of land for economic benefits. The key influence factors of spatial variation in land use are altitude; gradient; and the distance to rivers, highways and villages. (3) A plan of tourism land management and control should be established with construction land as the key indicator.

## 1. Introduction

As an important component of terrestrial ecosystems, mountain areas play an important role in regional ecological safety, and their rich natural resources are crucial to the development of society and the economy ^[1]^. A wide range of mountains in China cover 46.11% of China’s land area, and mountains constitute the most important ecosystem in China ^[2]^. Therefore, the exploitation and ecological construction of mountain resources are at the top of the list for sustainable development in China ^[3]^. With the rise and development of China’s tourism industry, mountain tourism has also developed rapidly, but the rise of mountain tourism has intensified its impact on the environment. While mountain tourism promotes local GDP growth and employment, and promotes the development of related industries, it also leads to changes in land use structure and natural ecological balance. For example, the construction of architectural facilities in scenic spots may ruin the inherent natural ecology of the local area and break the balance of natural land use. New trails and activities of tourists inevitably affect the ecosystem and biodiversity. The development of mountain tourism has led to the continuous encroachment and encroachment of the effective habitats of species in nature reserves ^[4-7]^, and the fragmentation of habitat patches constitutes the greatest threat to mountain biodiversity ^[8-10]^.In China more than 22% of the natural reserve area has been ruined due to unscientific tourism development, and 11% has shown resource degradation ^[11,12]^. Land is the basis for tourism development, and exploiting tourism resources and constructing tourism infrastructure can change the surface structure, land, soil and vegetation, and the corel change is the land use. Therefore, the research on the spatial changes and influencing influencing factors of tourism land in mountainous areas constitutes an important part of the research on the environmental impact of mountain tourism.

In researching the spatial variation in tourism land and the driving factors, scholars in China have employed geographic information system (GIS) and remote sensing (RS) technologies to interpret and compare two or more RS images to assess the changing trends of mountain tourism land variation. It is commonly agreed that with the development of the tourism industry, land use has undergone an increase in scenic spots, transportation and infrastructure and a decrease in farm land and ecosystem. Natural environmental experiences tend to cause changes. However, in different areas, the features and trends of changes in tourism land vary ^[13]^. Research on the Lushan Nature Reserve shows that the vegetation is undergoing positive secondary changes. The fragmentation level of vegetation in the outer area is significantly higher than that in the inner area. The total amount of arable land, woodland and unused land in the reserve has decreased in the past 15 years, while garden plots, grasslands, urban villages, industrial and mining land, transportation land, water area and water conservancy facilities land has increased significantly ^[14]^. Research on Yangshuo County, a famous mountain resort, shows that with the development of tourism, the scale of tourism land continues to increase, the area of arable land generally decreases, while other non-agricultural land increases rapidly ^[15]^. The study of Emeishan City found that the use of tourist land presents a spatial expansion pattern from scenic spots to urban areas along the passage, and the growth of tourist land in protected areas tend to increase without restrictions ^[16]^. Some research also found that after tourism development, mountainous land use types of rural tourism destinations are diversified, leading to complicated land functions ^[17-18]^. In terms of influencing factors, it is generally believed that the factors leading to the change of tourist land in mountainous areas are diverse, mainly including the increase in the area of transportation facilities and tourist facilities caused by the development of tourism ^[13-14,19]^. In addition to tourism, other factors, such as government decisions ^[20]^,demographic changes ^[21]^, natural factors, rapid urbanization^[14]^, the coercive effect of residents on forest parks ^[22]^, feedback from weakening agricultural development, feedback from weakening industrial development ^[19]^, farmers abandoning agriculture for business, the increase of urban and village and industrial and mining land ^[15]^ may also lead to changes in land use in mountain tourism. In addition, some scholars believe that the pioneering farmers in rural tourism have played a leading role in the evolution of land use ^[23]^. The change of tourism land, based on and driven by the internal driving factors of the tourism industry, with the external driving factors as supporting forces, results from both internal and external factors ^[24-26]^.

The research on tourist land in mountainous areas in foreign countries predates China and some results have been achieved. Scholars use multi-temporal land satellite images ^[27-30]^?Spatial analysis ^[31-32]^, patch analysis ^[33]^ and other methods are used to analyze the land use changes and influencing factors in the case tourism area. It is believed that in addition to the rapid development of tourism, global climate change and regional policies have also made certain contributions to changes in land use types and changes including habitat loss and landscape fragmentation. Ecological perspective is also one of the methods commonly used by scholars to study mountain tourism land. For example, Kurniawan has studied the change patterns of landscape in mountainous islands, and found that land use/land cover tends to change based on the distance of routes, ports, coastlines, public services, rural centers, business districts, settlements, tourist accommodation, tourist centers, tourist attractions, and and landfill sites ^[34]^. Another example, Mwalusepo’s case study of the Unguja tourist area on Zanzibar Island in Tanzania shows that tourism development, climate change, land use and land cover changes have had an important impact on human communities and human ecosystems ^[35]^. Some scholars have also studied from the perspective of disaster science. For example, Jaydip’s research on Uttarakhand in India found that with the development of mountain tourism, the land use is expanding from gentle slopes to steep slopes, and construction on steep slopes may easily cause natural disasters such as landslides ^[30]^. Beautiful landscapes are the material basis for the development of rural tourism, some scholars emphasize that the development and management of mountain tourism should pay attention to the analysis of the suitability of land use ^[36]^.

This research is inspired by the above achievements,but problems still exist as follows. First, the existing documents are mainly the analyses of specific case scenic spots and research on multiple scenic spots in a certain area is relatively lacking. The analysis of a single scenic spot only reflects the land use change of individual scenic spots, while the study on multiple scenic spots in a region reflects the characteristics of the entire region. Second, as for the analysis of influencing factors, model quantitative analysis is insufficient. Most of the previous studies focused on simple quantitative description. Third, from the perspective of research, most of the previous studies have focused on the impact of human activities on land use, and little was done from the physical and geographic perspectives such as mountain height, direction, and slope. This paper’s object is Luanchuan County in central China, which features a small population, rapid tourism development, a large share of the tourism industry and many scenic spots. This study employs methods such as land use data analysis, field microeconomic surveys, and logistic regression analysis to study tourism land spatial variation in mountain areas and influence factors to enrich the related literature.

## 2. The Studied Areas, Data Sources, and Methods of Research

### 2.1 The Studied Areas

Luanchuan County, lies in the southwest part of Henan Province with the geographic coordinates 111°11′∼112°01′E, 33°39′∼34°11′N and is bordered by the Funiu Mountains at the southwest corner of Luoyang city (Picture 1). Luanchuan County covers a total area of approximately 2478 km^2^ and contains 14 townships and 209 administrative villages with a total population of approximately 350,000 according to Luoyang City Statistics Bureau^[37]^. Luanchuan County is 150 km from Luoyang city and 300 km from the capital of Henan Province, Zhengzhou city; provincial and national highways intersect there, making transportation easy. It is a classic mountain area with mountains of varying height. The highest elevation is 2212.5 m, and the lowest elevation is 450 m, with an altitude difference of 1762.5 m for a typical undulating mountain topography. It is the highest county in Henan Province, and its urban area is 750 m above sea level. Luanchuan County has many rich and high-ranking tourism resources. There are 8 main classes, 26 subclasses and 84 primary classes of sightseeing resources, constituting 54.2% of the national tourism resources. Luanchuan County boasts two 5A scenic spots, five 4A scenic spots, and thirteen A-level sightseeing zones (Picture 1). The tourism industry is the key industry in Luanchuan County.

### 2.2 Data Sources and Processing

Data on land use and variation were collected from the first and second land use status surveys by the local government in 1991 and 2010, respectively. Data on land use in 2018 were collected by the author on the basis of a Google satellite field investigation. First, 0.6 m high-resolution satellite map tile data were downloaded from MapDown and used to form a satellite map of Luanchuan County. Then, land use data for 2001 were collected by remote sensing image data in the study area, land use data for 2018 were collected by Erdas, and land use distribution data were collected by field investigation for comparison.

The effect factors of land use changes are based on the author’s analysis of the sampling point map. In view of the centrality of land use conversion, this paper focuses on four types of conversion, namely, woodland to farmland, woodland to construction land, woodland to water areas, and farmland to construction land; these four types of conversion constitute 81.25% of the land area. This paper employs balanced stratified sampling to choose sampling points and controls the number of sampling points over 2000 to make the research representative. After the sampling points were chosen, type maps of land use changes were overlapped with distribution maps of topography, slope direction, villages and rivers, and the attributes of the sampling points were read manually.

### 2.3 Methods of Research

#### 2.3.1 Plane Divergence Rate

The plane divergence rate, usually studied in association with the land use expansion rate (dynamic degree), refers to the expansion speed of different types of land in different research periods in the same area^[38]^.

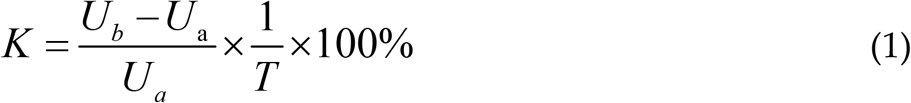

As indicated in Formula 1, K represents the land use divergence rate in the research period; *U*_*b*_ and *U*_*a*_ refer to the portions of tourism land at the beginning and end of the research period, respectively; and *T* is the length of the research. When *T* is set as one year, *K* is the annual variation rate of land use in Luanchuan County.

#### 2.3.2 Logistic Regression Analysis

This paper analyses geographic space factors that affect tourism land by multiple logistic regression. Assuming that X is a response variable and P is the response probability of the model, then the regression model can be given as follows:

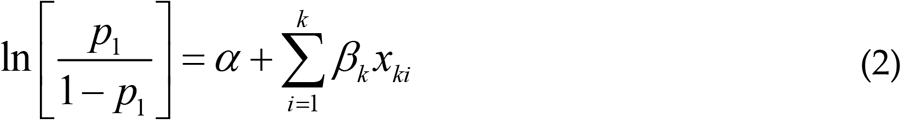

In Formula 2, *P*_*1*_*=P (y*_*i*_*=x*_*1i*_,*x*_*2i*_,*…,x*_*ki*_) is the occurrence rate of the event in the given series of independent variables *x*_*1i*_,*x*_*2i*_,*…, x*_*ki*_, with *β* as slope and *α* as intercept. For the actual calculation, the logistic function of SPSS16.0 statistical software is recommended.

#### 2.3.3 Classification of tourist land types

In view of the lack of a complete classification system for domestic tourism land, for the convenience of research, this article adheres to the principle of combining the natural attributes and use attributes of the land, and the universality and particularity, referring to Su Kun, Zhou Yong ^[39]^, Wang Jinye ^[40]^, Lu Weimin, Liu Yang ^[41]^, Peng Hui, Bi Yuzhu ^[42]^, etc., divided the land types of scenic spots within the study area. Tourism land mainly includes 5 first-class categories (tourism construction land, tourist agricultural land, tourist forest land, tourist water body land, tourist unused land), 9 second-class categories (land for scenery viewing and recreation, tourist facilities and engineering facilities, Land for tourism and commercial service facilities, land for management services, residential and social comprehensive land, agricultural land, water body land, unused land), 18 three-level categories (land for natural landscape viewing, landscape green space, cultural landscape viewing land, tourist facility land, engineering Facilities land, tourism shopping land, tourism catering land, tourism accommodation land tourism and entertainment land, management service land, residential area, integrated service land, cultivated land, garden land, water body land, unused land, etc.).

## 3. Tourism Land Spatial Variation

Over the past 28 years (1991-2018), both the number of scenic spots and the area of tourism land have increased remarkably. The number of scenic spots increased from 1 to 13, and the area of tourism land grew from 2.11 km^2^ to 159.69 km^2^, a 74.7-fold increase. With the expansion of the scale of tourism land use, the spatial structure of tourism land experienced great changes.

### 3.1 Significant Tourism Land Expansion in Scenic Spots

Tourism resource development and scenic spot construction include the construction and operation of various infrastructure and production facilities, namely, the spatial expansion of construction land. In the early stage of the development of scenic spots, there are a small number of necessary facilities, such as roads, gates, service centres, restaurants, cable cars, and cableways. When scenic spots become famous and experience a surge of visitors, the old facilities can no longer satisfy consumers’ needs or support the expansion of related facilities or the development of farmhouse resorts, restaurants and new scenic spots. In addition to construction projects, the construction of landscape and afforested areas are high priorities.

The plane divergence rate of internal construction in scenic spots is proportional to the location and ranking of scenic spots. As shown in Table 1, all 13 scenic spots show an increase in area, while their speed of change differs. Scenic spots with higher rankings near major towns, such as Laojun Mountain, Longyuwan National Forest Park, Hudie Valley, the scenic Yangzigou region, and the ski resort, have developed rapidly. Lower-ranked scenic spots far from major towns and highways, such as the Chongdugou scenic spot and the Hongdoushan scenic spot, have developed more slowly.

**Table 1.** Scenic spot construction land expansion at the beginning and end of the study.

| Scenic spot | Area at the beginning of the research (hm<sup>2</sup>) | Area at the end of the research (hm<sup>2</sup>) | Expanded area (hm<sup>2</sup>) | Dynamic degree (%) |
| --- | --- | --- | --- | --- |
| Chongdugou scenic spot | 29.98 | 48.66 | 18.68 | 62.29 |
| Yangzigou scenic spot | 28.42 | 48.22 | 19.81 | 69.7 |
| Longyuwan National Forest Park | 47.3 | 92.13 | 44.82 | 94.76 |
| Laojun Mountain | 34.39 | 60.42 | 26.02 | 75.67 |
| Jiguandong | 6.69 | 8.35 | 1.65 | 24.71 |
| Ski Resort | 5.99 | 92.13 | 32.2 | 53.75 |
| Jiulong Mountain | 2.29 | 2.32 | 0.02 | 1.01 |
| Daohuigou | 28.92 | 34.85 | 5.93 | 20.52 |
| Baoduzhai | 39.15 | 41.17 | 2.02 | 5.17 |
| Hudie Valley | 2.98 | 34.8 | 31.82 | 1.12 |
| Hongdoushan scenic spot | 6.87 | 6.93 | 0.07 | 0.98 |
| Great Canyon rafting | 15.22 | 15.79 | 0.58 | 3.82 |
| Chongdugou scenic spot rafting | 14.39 | 14.61 | 0.22 | 1.56 |

In addition, the expansion rate of scenic spot construction land is related to the level and nature of development. Construction land expands at high speed in the beginning during development in scenic spots, while it slows down or even stops in well-developed scenic spots because of adequate facilities. Some scenic spots, such as Jiulong Mountain Springs, are unlikely to expand rapidly due to restricted resources and the influence of nature. The Jiulong Mountain Spring resources are exploited to the limit, so it is impossible and unnecessary to expand in depth. Jiguandong scenic spot, featuring karst caves, has reached maximum development and requires no more construction land. For two rafting scenic spots, no further construction projects are needed, so there is little increase in construction land.

### 3.2 Diversified Functions of Scenic Spots Land

With the intensive development of tourism, the land use types of scenic spots change from single to compound and diversified, and the number of land use types also increases. As the main land use type of scenic spots, construction land becomes more diversified in its functions. At the beginning of the development of some scenic spots, the types of construction land that cater to sightseeing and accommodation are simple. When scenic spots are developed, the types of construction land increase to meet most tourism needs. Investigation shows that among 13 scenic spots in Luanchuan County, most appear to be diversified in land use types, except for single-function spots such as ski areas, Great Canyon rafting, and Chongdugou scenic spot. Laojun Mountain has a planned high starting level, and the Hongdoushan scenic spot has lagging development (Table 2). At the classic Chongdugou scenic spot, the residential areas, farmland, woodland and water areas are being converted to various types of tourism land. Traditional farmland is used to cultivate fruits, herbs, flowers, etc. Construction land originally designated for roads, urbanization, and residential areas is transformed into commercial tourist services land (accommodation, restaurants and recreational facilities), tourism production land, and infrastructure land. In short, in the tourism development process, the diversified service functions of scenic spots become increasingly prominent. The land service function develops from residence and production to entertainment.

**Table 2.**
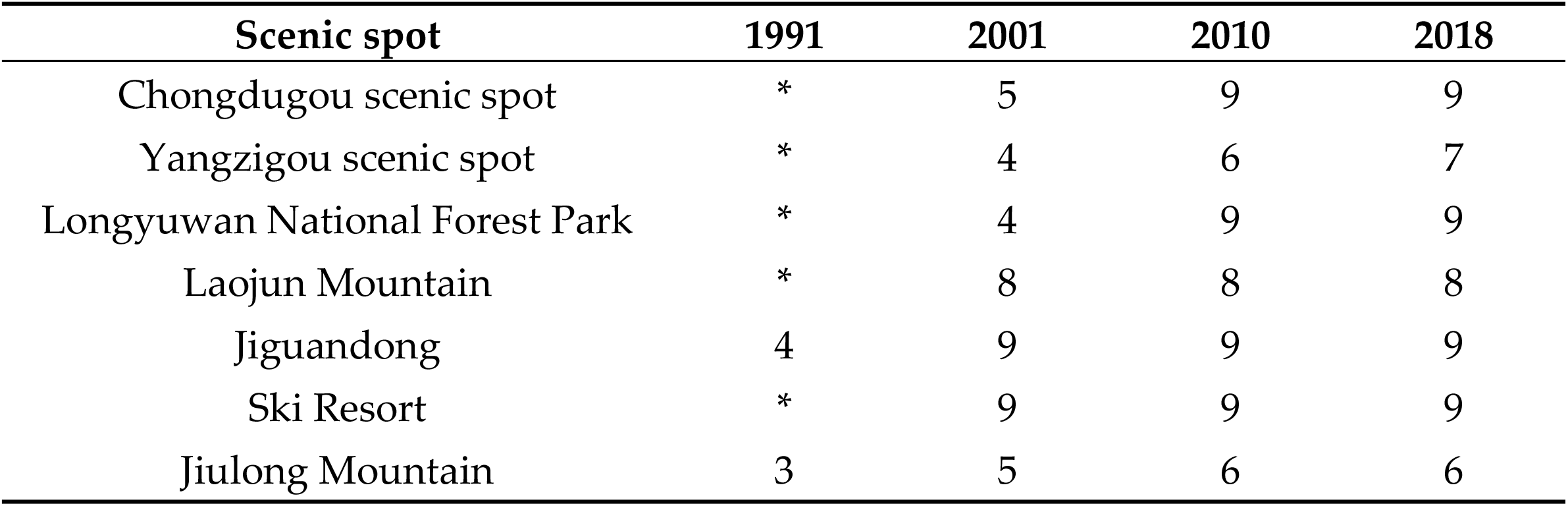
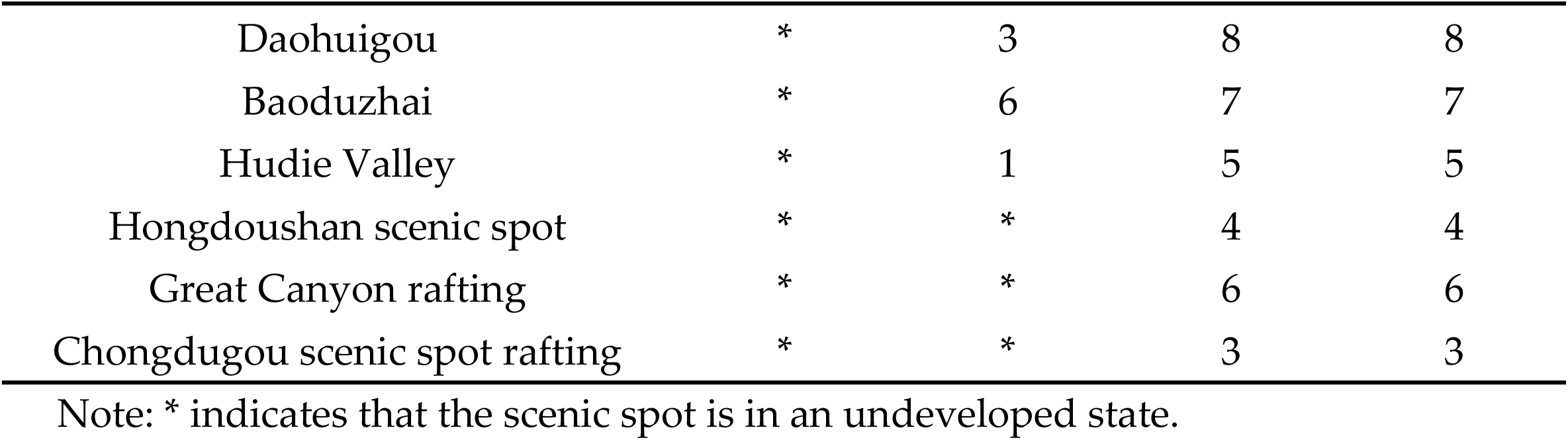
Changes in types of tourism land in various scenic spots

| Scenic spot | 1991 | 2001 | 2010 | 2018 |
| --- | --- | --- | --- | --- |
| Chongdugou scenic spot | * | 5 | 9 | 9 |
| Yangzigou scenic spot | * | 4 | 6 | 7 |
| Longyuwan National Forest Park | * | 4 | 9 | 9 |
| Laojun Mountain | * | 8 | 8 | 8 |
| Jiguandong | 4 | 9 | 9 | 9 |
| Ski Resort | * | 9 | 9 | 9 |
| Jiulong Mountain | 3 | 5 | 6 | 6 |
| Daohuigou | * | 3 | 8 | 8 |
| Baoduzhai | * | 6 | 7 | 7 |
| Hudie Valley | * | 1 | 5 | 5 |
| Hongdoushan scenic spot | * | * | 4 | 4 |
| Great Canyon rafting | * | * | 6 | 6 |
| Chongdugou scenic spot rafting | * | * | 3 | 3 |

Note: \* indicates that the scenic spot is in an undeveloped state.

Considering the lack of a complete classification system of domestic tourism land, to facilitate the research and classify tourism land use types in Luanchuan County, this paper uses the principle of combining natural land attributes with use attributes and generality with particularity, referring to the achievements of Su, K. and Zhou, Y. (2008) ^[17]^, Wang, J., et al. (2015) ^[18]^, Lu, W. and Liu, Y. (2016) ^[19]^, and Peng, H. and Bi, Y. (2015) ^[20]^. Land for tourism use is classified in 5 main categories (tourism construction land, tourism farmland, tourism woodland, tourism water areas, and tourism untouched land), 8 secondary categories (sightseeing and vacation land, sightseeing facilities and project construction land, commercial tourist service facilities land, management land, village community land, farmland, water areas, and untouched land), and 16 tertiary categories (sightseeing land for natural landscape, green space, sightseeing land for human landscape, sightseeing facilities land, project facilities land, shopping land, restaurant land, accommodation land, recreational land, management land, residential land, comprehensive services land, farmland, gardens, water areas, and untouched land).

### 3.3 Rapid Transformation of Land Use Types

As the tourism industry expands and the number of scenic spots increases, tourism land areas undergo a significant increase, and land use types in scenic spots change greatly. During 1991-2018, the structural changes in tourism land in Luanchuan County featured a sharp increase in construction land, a slight increase in water areas and a decrease in farmland. From the beginning period to the development period (1991-2010), the main types of land in Luanchuan County scenic spots were woodland and farmland, which together made up over 90% of the total. The development of the tourism industry has not affected local people’s traditional life and production style. Some land use types in scenic spots, especially farmland, are used mainly to satisfy local needs and are not completely taken over by scenic spots. Locals in scenic spots still pursue agriculture and forestry as their main ways to make a living and tourism as a minor way. The tourism industry reached a certain scale in the development period to the transformation period, 2010-2018 (except for two rafting scenic spots built in 2010). Consequently, all land in scenic spots is used to develop tourism, and many types of land are changing to construction land. A small amount of remaining farmland serves mainly as picking gardens or to cultivate flowers. Luanchuan County has been following an economic strategy of “a rich county of tourism” since 2000, so water areas have increased gradually due to a series of policies designed to protect woodlands and expand water areas (Table 3).

**Table 3.** Luanchuan County tourism land areas during 1991-2018 (hm^2^)

| Land use type | 1991 | 2001 | 2010 | 2018 |
| --- | --- | --- | --- | --- |
| Construction land | 8.98 | 358.97 | 390.64 | 411.78 |
| Farmland | 4.57 | 689.39 | 218.78 | 133.07 |
| Woodland | 186.38 | 13809.06 | 15100 | 15217.12 |
| Water area | 1.04 | 169.54 | 152.04 | 158.49 |
| Untouched land | 10.13 | 59.77 | 115.61 | 90.56 |
| Total | 211.1 | 15086.73 | 15977.07 | 16011.02 |

**Table 4.**
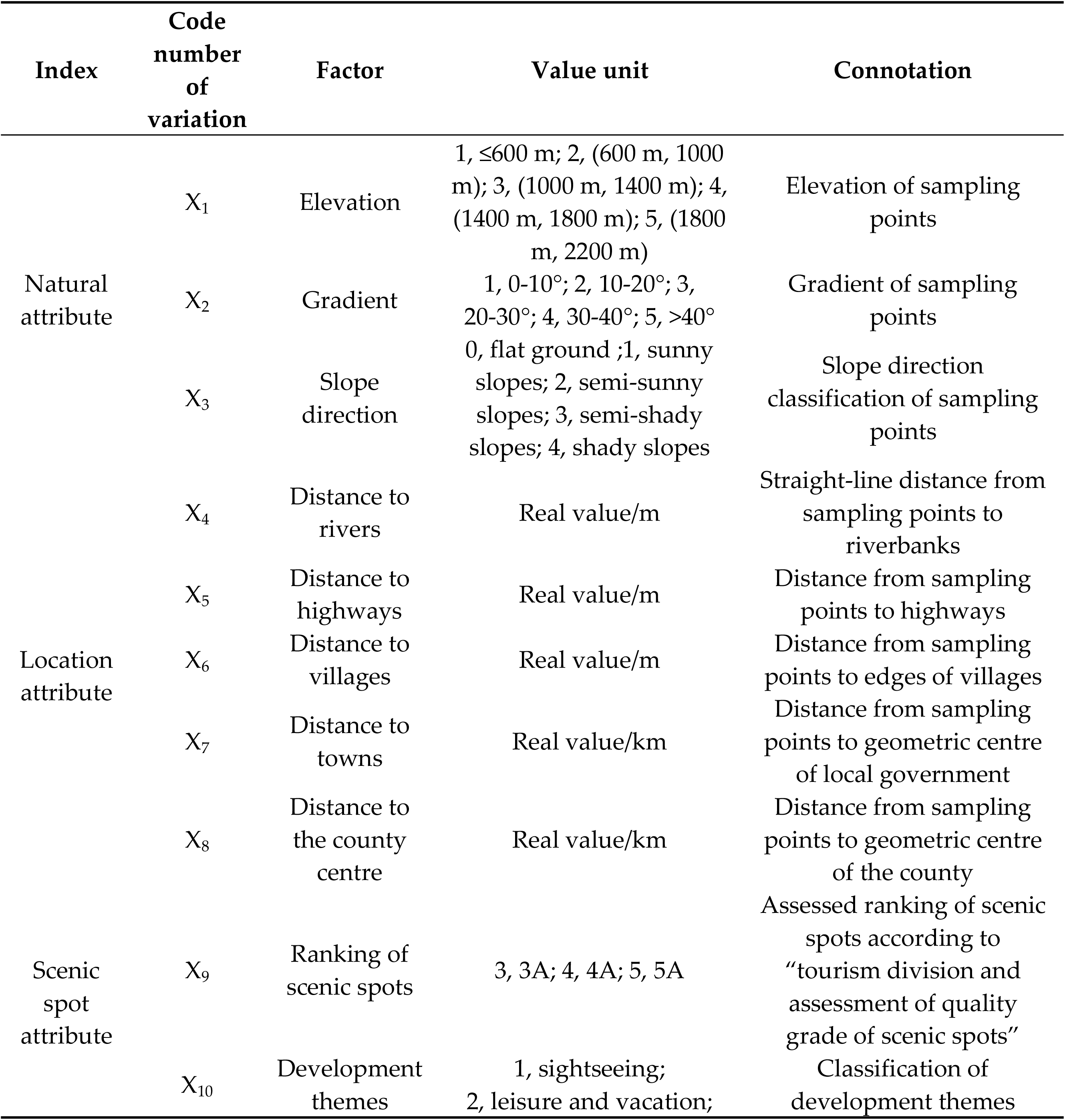
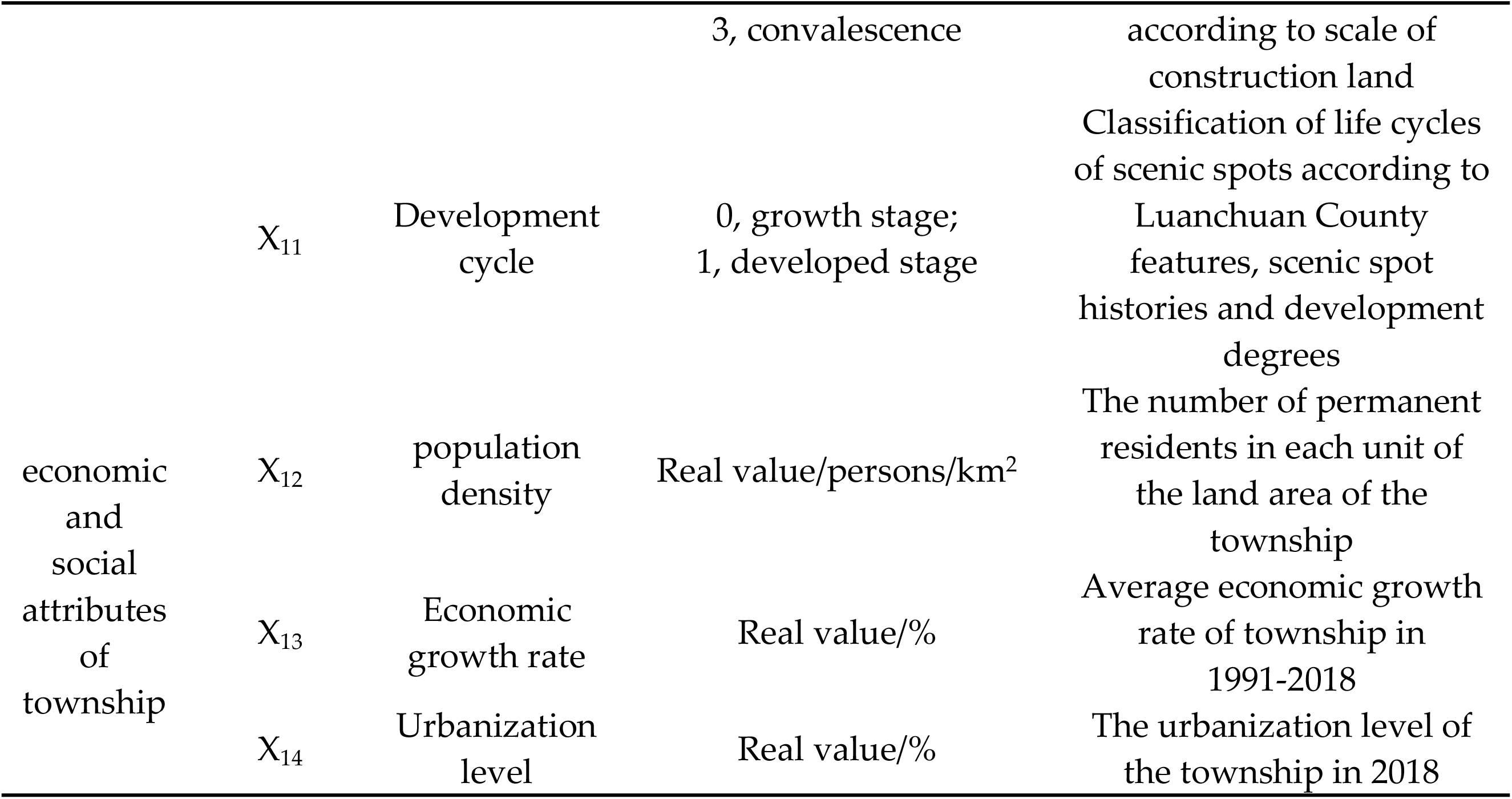
Variation designs

## 4. Influence Factors of Tourism Land Spatial Variation

### 4.1 Variation Designs

The changes in tourism land use types result from human utilization and development of land for economic benefits. In the short term, land use type changes outside the natural environment make no difference. If people crave a greater economic return through utilization and development, they must take account of the natural factor of the land, which directly determines the cost and profits of exploitation. Therefore, in regard to various designs, the altitude, gradient and direction affect tourism land changes. To secure better earnings, people must also consider the location factors. Because there are great differences in development necessity and earnings between different locations, land utilization is not absolutely balanced in space. Some areas are worth developing (economically and technically) and can be developed, while other places are not. Therefore, the influencing factors of land use types include distances to villages, roads, rivers and centres of a certain area. Different types of scenic spots have different construction land features and development histories, so the scales of land use and structures may vary greatly, reflecting a particular scenic spot’s construction demand. As a result, attributes such as scenic spot rankings and development themes exert a strong influence on the transformation of land use types. Scenic spot exploitation is subject to regional economic and social conditions. Different regional conditions and features affect the importance attached to scenic spot exploitation, policies, investment and necessity of development. Therefore, population density, speed of economic growth and the level of urbanization of the towns where scenic spots are located might affect the transformation of land use types. This paper analyses the influence of 14 factors and four types of indexes: natural attribute, location attribute, scenic spot attribute and township social attribute.

### 4.2 Analysis of Influence Factors of Main Transformation Types

At the first level of tourism land, there are five land types. So, there should be 20 types of any two types of mutual conversion, but the actual number of conversion types cannot reach the upper limit. The common four types of land conversion areas have been accounted for 76.8% of the total conversion area. Therefore, considering the limitation of paper space, it mainly analyzes four types of land conversion.

#### 4.2.1 Farmland to Construction Land

A value of 1 is used for land use transformation types caused by variations, such as “farmland to construction land”, and 0 is used for other types. The logistic analysis results shown in Table 2 indicate that factors such as altitude, gradient, distance to rivers, highways, and distance to villages are significant (Table 5, Model 1).

**Table 5.**
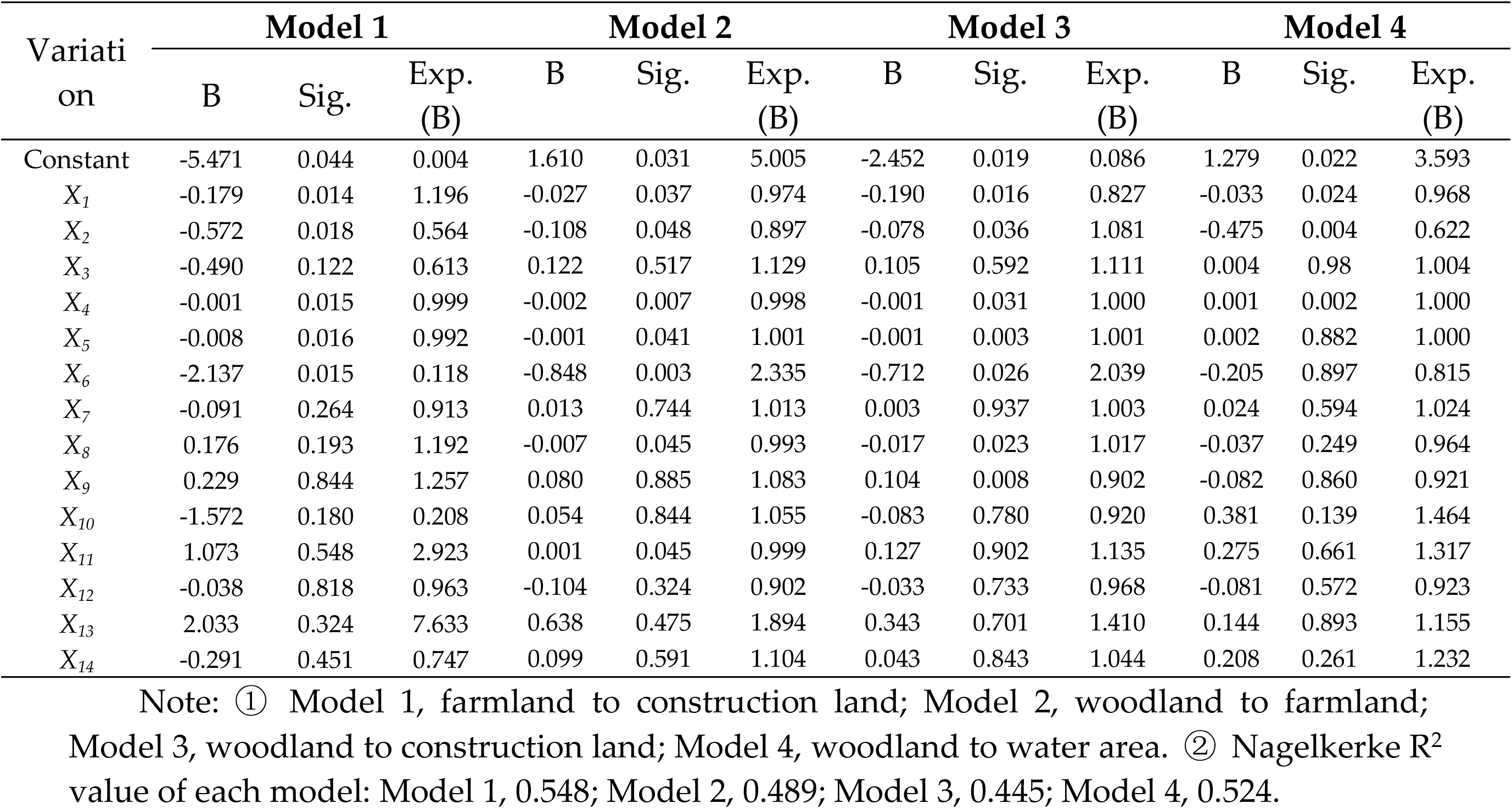
Key indicators of model analysis results

| Variation | Model 1 |  |  | Model 2 |  |  | Model 3 |  |  | Model 4 |  |  |
| --- | --- | --- | --- | --- | --- | --- | --- | --- | --- | --- | --- | --- |
|  | B | Sig. | Exp. (B) | B | Sig. | Exp. (B) | B | Sig. | Exp. (B) | B | Sig. | Exp. (B) |
| Constant | -5.471 | 0.044 | 0.004 | 1.610 | 0.031 | 5.005 | -2.452 | 0.019 | 0.086 | 1.279 | 0.022 | 3.593 |
| $X_1$ | -0.179 | 0.014 | 1.196 | -0.027 | 0.037 | 0.974 | -0.190 | 0.016 | 0.827 | -0.033 | 0.024 | 0.968 |
| $X_2$ | -0.572 | 0.018 | 0.564 | -0.108 | 0.048 | 0.897 | -0.078 | 0.036 | 1.081 | -0.475 | 0.004 | 0.622 |
| $X_3$ | -0.490 | 0.122 | 0.613 | 0.122 | 0.517 | 1.129 | 0.105 | 0.592 | 1.111 | 0.004 | 0.98 | 1.004 |
| $X_4$ | -0.001 | 0.015 | 0.999 | -0.002 | 0.007 | 0.998 | -0.001 | 0.031 | 1.000 | 0.001 | 0.002 | 1.000 |
| $X_5$ | -0.008 | 0.016 | 0.992 | -0.001 | 0.041 | 1.001 | -0.001 | 0.003 | 1.001 | 0.002 | 0.882 | 1.000 |
| $X_6$ | -2.137 | 0.015 | 0.118 | -0.848 | 0.003 | 2.335 | -0.712 | 0.026 | 2.039 | -0.205 | 0.897 | 0.815 |
| $X_7$ | -0.091 | 0.264 | 0.913 | 0.013 | 0.744 | 1.013 | 0.003 | 0.937 | 1.003 | 0.024 | 0.594 | 1.024 |
| $X_8$ | 0.176 | 0.193 | 1.192 | -0.007 | 0.045 | 0.993 | -0.017 | 0.023 | 1.017 | -0.037 | 0.249 | 0.964 |
| $X_9$ | 0.229 | 0.844 | 1.257 | 0.080 | 0.885 | 1.083 | 0.104 | 0.008 | 0.902 | -0.082 | 0.860 | 0.921 |
| $X_{10}$ | -1.572 | 0.180 | 0.208 | 0.054 | 0.844 | 1.055 | -0.083 | 0.780 | 0.920 | 0.381 | 0.139 | 1.464 |
| $X_{11}$ | 1.073 | 0.548 | 2.923 | 0.001 | 0.045 | 0.999 | 0.127 | 0.902 | 1.135 | 0.275 | 0.661 | 1.317 |
| $X_{12}$ | -0.038 | 0.818 | 0.963 | -0.104 | 0.324 | 0.902 | -0.033 | 0.733 | 0.968 | -0.081 | 0.572 | 0.923 |
| $X_{13}$ | 2.033 | 0.324 | 7.633 | 0.638 | 0.475 | 1.894 | 0.343 | 0.701 | 1.410 | 0.144 | 0.893 | 1.155 |
| $X_{14}$ | -0.291 | 0.451 | 0.747 | 0.099 | 0.591 | 1.104 | 0.043 | 0.843 | 1.044 | 0.208 | 0.261 | 1.232 |

Note: ① Model 1, farmland to construction land; Model 2, woodland to farmland; Model 3, woodland to construction land; Model 4, woodland to water area. ② Nagelkerke R<sup>2</sup> value of each model: Model 1, 0.548; Model 2, 0.489; Model 3, 0.445; Model 4, 0.524.

- Altitude Farmland, such as cultivated land, which spreads over the flats along riverbanks, is converted to tourism villages, such as farmhouse resorts. Farmland at low altitudes is adjacent to traffic routes, so traffic facilities are spread around it. The land use transformation type mostly seen in mountain areas is farmland to construction land, which means farmland area decreases as it is used to accommodate infrastructure for the development of tourism.
- Gradient Owing to cost and technological issues, common construction projects are not usually situated on steep slopes. Therefore, gradient is a main influence factor when farmland is transformed into construction land. The infrastructure of scenic spots and tourism villages is built on farmland with a slight slope, on river terraces, or near rivers, highways and villages. Gradient is also considered when choosing a tourism route.
- Distance to rivers Rivers are an important water source for human survival and development. In mountain areas, the distribution of the population is consistent with that of rivers that run through the lower-elevation areas. The farmland along rivers is flat or has slight slopes. Therefore, the distance to rivers is related to the distribution of population, residential areas and farmland. During the scenic spot development process, the tourism infrastructure, such as farmhouse resorts, hotels, restaurants, and shops, tends to be situated near rivers and spreads along riverbanks. The farther a site is from a river, the more unlikely it is that tourism infrastructure will be built there. In mountain areas, tourism routes are built along rivers because rivers flow through flat areas with a longitudinal gradient.
- Distance to highways In mountain areas, the main roads follow the course of rivers, so distance to highways has the same influence as distance to rivers. To build main roads in mountain areas, technology, cost and safety must be considered. The roads must follow a gentle slope and require small investment and easy construction, and riverbanks meet these criteria. Therefore, the direction of roads is relative to that of rivers. During the tourism development process, the transformation of farmland to construction land is influenced by distance to highways. The nearer a site is to a highway, the more possible this transformation is. The farther a site is from a highway, the more impossible this transformation is.
- Distance to villages In mountain areas, the location of villages is closely related to the natural environment, and the distribution of villages is an outcome of historical development. In the formation and developing processes of villages, the natural environment must be considered. During the process of transforming farmland to construction land, tourism infrastructure tends to be built in or near the existing villages. Therefore, distance to villages is a crucial factor in transforming farmland to construction land. Many villages transform and rebuild houses in original residential areas into farmhouse resorts. Some tourism infrastructure, such as farmhouse resorts, is built on farmland near villages.

#### 4.2.2 Woodland to Farmland

A value of 1 is used for land use transformation types caused by variations such as “woodland to farmland”, and 0 is used for other types. The logistic model results shown in Table 2 indicate that factors such as altitude; gradient; distance to rivers, highways, villages, and the county centre; and the scenic spot development cycle are significant (Table 2, Model 2).

The transformation of woodland to farmland in scenic spots is driven by that of farmland to construction land. In the tourism development process, farmland decreases because it is transformed to construction land. To compensate for the loss of farmland, woodlands that are not far from rivers, highways and villages and have a low relative elevation and gentle slopes are transformed to farmland. Therefore, factors such as distances to rivers, highways, villages; altitude; and gradient are deciding factors in transforming woodland to farmland.

In Model 2 (woodland to farmland), distance to the county centre is significant. As a regional centre, the county affects the exploitation of tourism resources. In Luanchuan County, scenic spots near the county centre have a long history of a high degree of mature exploitation on a large scale. These scenic spots have caused the loss of farmland, so most woodlands are converted to farmlands. The local industrial structure is improved by tourism development, but agriculture remains, and retaining farmland shows the feature of industrial diversification in Luanchuan County. In fact, the number of people engaged in tourism increases dramatically when the tourism industry expands. However, not all agricultural populations are engaged in the tertiary industry.

The period of scenic spot development has a strong influence on the transformation of woodland to farmland and shows the overall feature of scenic spot development, that is, there is a great difference between developing scenic spots and developed ones. After long-term development, the transformation of types of land use in developed scenic spots is complex and diverse, especially in terms of woodland to farmland. Fully developing scenic spots is the main reason for turning woodland into farmland. The formation mechanism is based on the supplementary mechanism of farmland decrease. As farmland is turned into construction land increasingly quickly, the evolution of the regional industrial structure requires compensating for farmland to maintain a balanced local economic and social structure.

#### 4.2.3 Woodland to Construction Land

A value of 1 is used for land use transformation types caused by variations such as “woodland to construction land”, and 0 is used for other types. The logistic model results shown in Table 2 indicate that factors such as altitude; gradient; distances to rivers, highways, villages, and the county centre; and scenic spot rankings are significant (Table 2, Model 3).

When more land is needed to construct scenic spots, and there is not enough farmland, woodlands that are located on gentle slopes with low altitudes and near rivers, villages and highways are transformed directly to construction land. Therefore, the above factors reach a high level. The nearer the county centre and the more developed the scenic spots, the more land they cover and the more decisive the factor is. Rankings of scenic spots represent the operational scale and level of development. High rankings indicate higher development levels, larger areas and a greater possibility of woodland being converted to construction land.

#### 4.2.4 Woodland to Water Area

A value of 1 is used for land use transformation types caused by variations such as “woodland to water area”, and 0 is used for other types. The logistic analysis results of Model 4 shown in Table 2 indicate that factors such as altitude, gradient, and distance to rivers are significant (Table 2, Model 4).

Woodlands are transformed to water areas by damming to store water. To build landscapes and restore the natural ecology, dams are built to store water with the original watercourse at the centre, or a reservoir is built on low terrain. Thus, woodlands are transformed into water areas. Factors such as altitude, gradient, and distance to rivers are significant. This transformation is rare in natural environments but constitutes a large proportion of land use type conversion.

## 5. Conclusion and Discussion

### 5.1 Conclusion

The rapid development of mountain tourism has caused drastic changes in the structure of land use and has had an important impact on the terrestrial ecosystem. Based on the land use data of four periods in the past 30 years, namely 1991, 2001, 2010, and 2018, with the land use expansion index and the spatial binary logistic adopted, research is made to study the spatial changes and influencing factors of mountain tourism land and the following conclusions can be obtained.

1. Spatial variation in land use in scenic spots shows the chain reaction of land use type transformation caused by construction land expansion, a key driving force of spatial variation in land use and land use type transformation. To satisfy the growing needs of tourism, more land is used to accommodate tourism facilities, resulting in the transformation of farmland to construction land, woodland to construction land, and woodland to farmland. Furthermore, woodlands are converted to water areas to improve the environment and build landscapes. In addition, with the intensive development of tourism, land use types of scenic spots change from single to compound and diversified, and the number of land use types also increases. As the main land use type of scenic spots, construction land becomes more diversified in its functions.
2. The changes in tourism land use types result from human utilization and development of land for economic benefits, and these changes are affected by many factors. The transformation of farmland to construction land is subject to altitude; gradient; and distances to rivers, highways, and villages. The transformation of woodland to farmland is affected by factors such as altitude; gradient; distances to rivers, highways, villages, and the county centre; and the scenic spot development cycle. The transformation of woodland to farmland is largely driven by that of farmland to construction land, and distance to the county centre and the scenic spot development cycle strongly influence land use. The transformation of woodland to construction land is greatly affected by factors such as altitude; gradient; distances to rivers, highways, villages, and the county centre; and scenic spot rankings. The transformation of woodland to water areas is affected by factors such as altitude, gradient, and distance to rivers.
3. A plan of tourism land management and control should be established with construction land as the key indicator. According to the above research, as for changes of tourism land use in mountainous areas, the change of construction land is the fundamental inducement, and construction land is the main driving mechanism leading to the change of various types of land use in the scenic area. Therefore, the planning and construction of tourist attractions should focus on the planning of tourism construction land. The construction land should match the development scale and development direction of the scenic spot, and should be appropriately controlled to be in harmony with the natural environment. Once the tourist attraction plan is determined, development and construction should be carried out in strict accordance with the plan to avoid blind expansion of construction land in the scenic spot, thereby avoiding disorderly changes in other land such as forest land and agricultural land. Although the occupation of construction land is inevitable for the development of tourism, the excessive spread of construction land is the main cause of the deterioration of the ecological environment of the scenic area. Therefore, it is of practical significance to establish a scenic environment management system with construction land as the core. Due to its particularity, the ecological environment of mountainous areas is fragile. When pursuing economic and social benefits from the development of mountainous tourist attractions, importance shoule alos be attached to improve ecological benefits in order to consolidate the foundation for sustainable development of mountain tourism. In addition, attention should be paid to coordinate the relationship between tourism planning and overall land use planning. The compilation of tourism planning should be based on the overall land use planning while the land use planning should fully consider the tourism land planning in the tourism planning, especially the tourism construction land planning.

### 5.2 Discussion

Forms of land use and spatial distribution changes are inevitable during the development process of mountain tourism. An analysis of the aspects of nature, location and economics is crucial to the understanding and control of land use changes of scenic spots in mountain regions. With Luanchuan County as an example, this paper analyses most of the scenic spots in a small area. However, considering the data accessibility and research period, the interspaces of the research period are not completely balanced. The types of land use transformation vary, but this analysis focuses on 4 of the most important types owing to space considerations. More types will be studied in the future. In addition, owing to workload considerations, when selecting samples of influence factors, this paper chose the minimum samples over 2000. In the future, a more precise portrait and analysis will be performed by increasing the number of samples.

## Author Contributions

Y. X. performed all the experiments and drafted the manuscript. All authors participated in the design of this study and analysis of results. Q. Z. conceived and coordinated this study.

## Acknowledgments

The authors would like to thank Academician Laboratory for Urban and Rural Spatial Data Mining of Henan Province for the support during the research. The authors would also like to thank AJE for providing linguistic assistance during the preparation of this manuscript.

---

**Figure 1.**
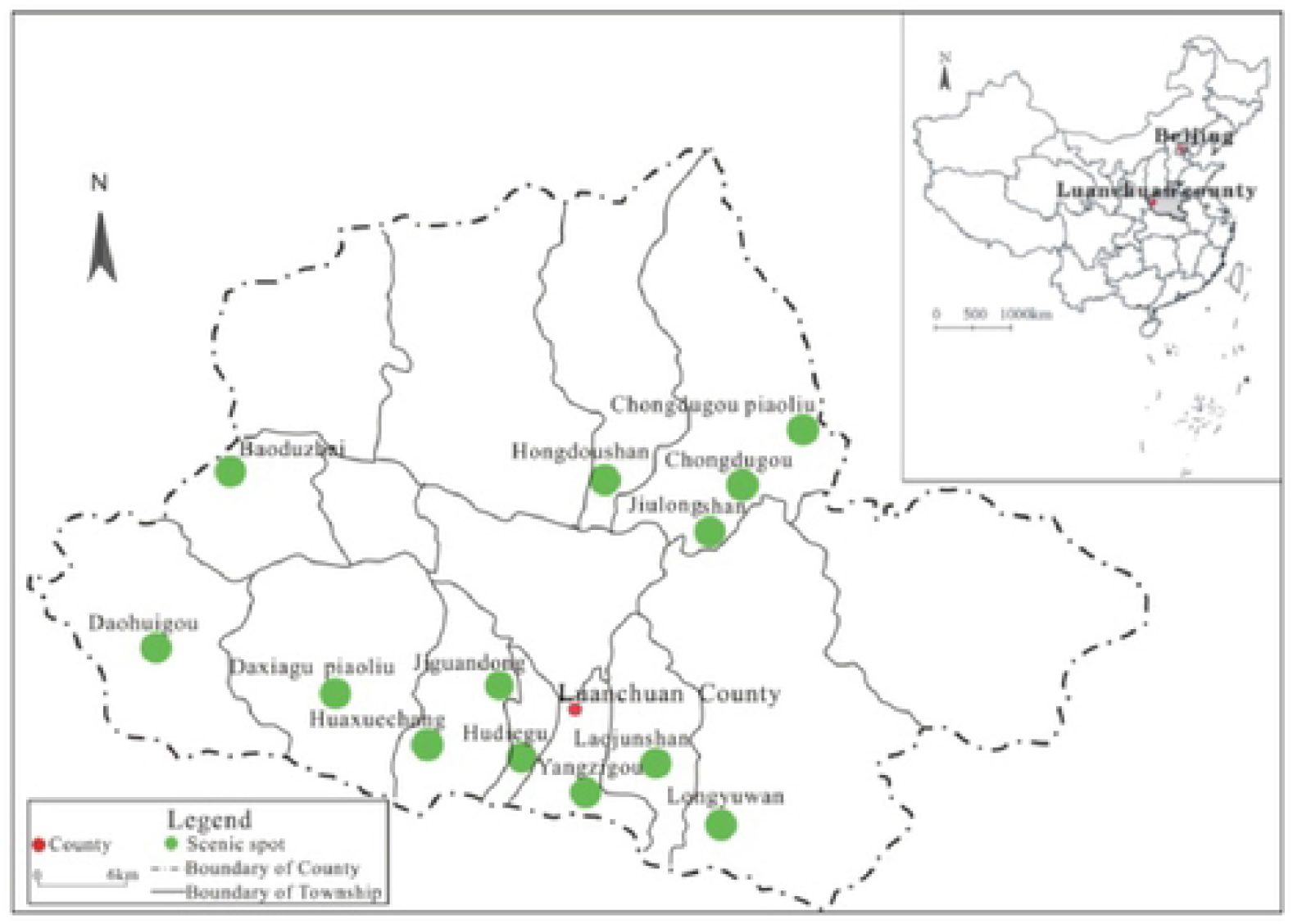
Distribution map of tourist attractions in Luanchuan County

